# Dog Savior: Immediate Scent-Detection of SARS-COV-2 by Trained Dogs

**DOI:** 10.1101/2020.06.17.158105

**Authors:** Omar Vesga, Andres F. Valencia, Alejandro Mira, Felipe Ossa, Esteban Ocampo, Maria Agudelo, Karl Čiuoderis, Laura Perez, Andres Cardona, Yudy Aguilar, Javier M. González, Juan C. Cataño, Yuli Agudelo, Juan P. Hernández-Ortiz, Jorge E. Osorio

## Abstract

Molecular tests for viral diagnostics are essential to confront the COVID-19 pandemic, but their production and distribution cannot satisfy the current high demand. Early identification of infected people and their contacts is the key to being able to isolate them and prevent the dissemination of the pathogen; unfortunately, most countries are unable to do this due to the lack of diagnostic tools. Dogs can identify, with a high rate of precision, unique odors of volatile organic compounds generated during an infection; as a result, dogs can diagnose infectious agents by smelling specimens and, sometimes, the body of an infected individual. We trained six dogs of three different breeds to detect SARS-CoV-2 in respiratory secretions of infected patients and evaluated their performance experimentally, comparing it against the gold standard (rRT-PCR). Here we show that viral detection takes one second per specimen. After scent-interrogating 9,200 samples, our six dogs achieved independently and as a group very high sensitivity, specificity, predictive values, accuracy, and likelihood ratio, with very narrow confidence intervals. The highest metric was the negative predictive value, indicating that with a disease prevalence of 7.6%, 99.9% of the specimens indicated as negative by the dogs did not carry the virus. These findings demonstrate that dogs could be useful to track viral infection in humans, allowing COVID-19 free people to return to work safely.

## Main

The only effective measure to ameliorate the impact of the COVID-19 pandemic is early and accurate identification of people infected with SARS-CoV-2. A key aspect of COVID-19 is that diagnostic tests must detect the virus in asymptomatic, pre-symptomatic and symptomatic patients^1,2^. The gold standard diagnostic test, the real-time reverse transcriptase-polymerase chain reaction (rRT-PCR), is not widely available in some countries due to poor reagent supply and low testing capacity^3^. Antibody tests are useless to prevent the dissemination of the virus, as they peak after the infectious period^4,5^. Social distancing and quarantines, as recommended by the WHO, have served to reduce the peak of infection and to provide “time to respond” for health authorities. While effective, they severely disrupt the economy. Balancing quarantine measures with the safe operation of the economy is a necessity. In particular, economic conditions of third world countries do not allow extended periods of quarantine. As the quarantine progresses, the risk of social disobedience increases, while the levels of poverty, malnutrition and violence, in the post-

COVID-19 era, could be more devastating than the pandemic itself. It is imperative to implement new, fast, and reliable diagnostic tests that offer novel opportunities for infection control and for safely re-opening the economy.

Humans have been using dogs - *Canis lupus familiaris* - for scent-detection since the beginnings of domestication^6^. Their sense of smell is powerful and useful; it is of no surprise that one of the first scientific attempts to study the olfactory capabilities of canines was published in *Nature* during the late 1880s^7^. Today, highly trained dogs are invaluable, as they are used to locate missing people, endangered species, explosives, narcotics, currency, criminals, cadavers, fossils, chemical pollutants, pests belonging to the arthropod and chordate phyla, and agricultural quarantine items, among others^8^. So far, analytical instruments have not surpassed the dog, mainly because in the vast majority of these specialized, complex and dangerous applications, perfect performance and complete reliability are imperative requirements: i.e. for the detection of landmines^9^.

Medical diagnosis is among the fields with potential usages for scent-detection by animals. Although this kind of research triggers hope and enthusiasm among journalists^10^, it receives little to no attention from practicing clinicians, who rely exclusively on semiology and sophisticated instruments to determine what afflicts their patients^11^. Previous efforts have reported the use of scent-specialized dogs to detect specific malignancies or infectious diseases, and dogs have been trained to alert their owners about the imminence of seizures, migraine crisis or glycemic changes. Unfortunately, these isolated studies often lack the scientific rigor necessary to validate a diagnostic test for clinical use^12^. However, some studies on scent-detection of pathogens by animals have demonstrated that with appropriate training and strict adherence to the scientific method, it is possible to obtain consistent results. For instance, trained dogs have been used to diagnose diseases related to the presence of the multi-drug resistant bacteria *Clostridiodes difficile*^13^, and trained giant African pouched rats (*Cricetomys gambianus*) have been used to detect pulmonary *Mycobacterium tuberculosis*^14^. Furthermore, a recent publication by the US Department of Agriculture (USDA) reported a comprehensive method to validate canine diagnosis of the plant pathogen *Candidatum* Liberibacter asiaticus. The authors demonstrated that detection of infected trees by dogs was superior to quantitative PCR (qPCR)^15^.

Dogs are able to detect and differentiate unique odors that result from the emission, from pathogens or tumors, of volatile organic compounds (VOC) that occur combined with the breath, respiratory secretions, saliva, feces, urine, skin, and/or sweat. This is the premise behind this approach: the “smell print” of the microorganism of interest^16^. In the case of SARS-CoV-2, a global effort centered at the Krogan laboratories at the University of California, San Francisco, revealed that 332 protein-protein interactions occur between the virus and the human patient during the infection process^17^. It is known that almost half of the VOC in the breath of normal humans contain nitrogen; thus, protein-protein interactions could lead to specific odors in the breath and respiratory secretions of COVID-19 patients that a dog could distinguished from the breath of healthy subjects^18^. In this work we designed and executed a dog training protocol that resulted in a reliable, cost effective and simple method for SARS-CoV-2 diagnostics.

### Scent-detection of SARS-CoV-2 in saliva of COVID-19 patients

We devised a safe device to put respiratory secretions obtained from SARS-CoV-2 positive patients in a sealed flask without exposing people or dogs to infection. The VOC are allowed to evaporate, so our dogs could smell them. We adapted the USDA validation method^15^, based on the experimental comparison of the dog performance against a gold standard (PCR in their study, rRT-PCR in this case), to determine the sensitivity (*SEN*), specificity (*SPC*), positive predictive value (*PPV*), negative predictive value (*NPV*), accuracy (*ACC*), and likelihood ratio (*LR*) of our dogs to diagnose COVID-19 *in vitro* (respiratory secretions) and *in vivo* (the body of the patient). We trained six dogs to identify the scent-mark of the pathogen with the highest possible accuracy: (1) Andromeda, intact female, 6 months old, a Belgian Malinois; (2) Nina, intact female, 25 months old, a Belgian Malinois; (3) Niño, castrated male, unknown age, an American Pit Bull Terrier; (4) Timo, intact male, 31 months old, a Belgian Malinois; (5) Vika, intact female, 36 months old, a Belgian Malinois; and (6) Vita, intact female, 36 months old, a first generation Alaskan Malamute x Siberian Husky (see **Fig. 1**).

**Figure 1:**
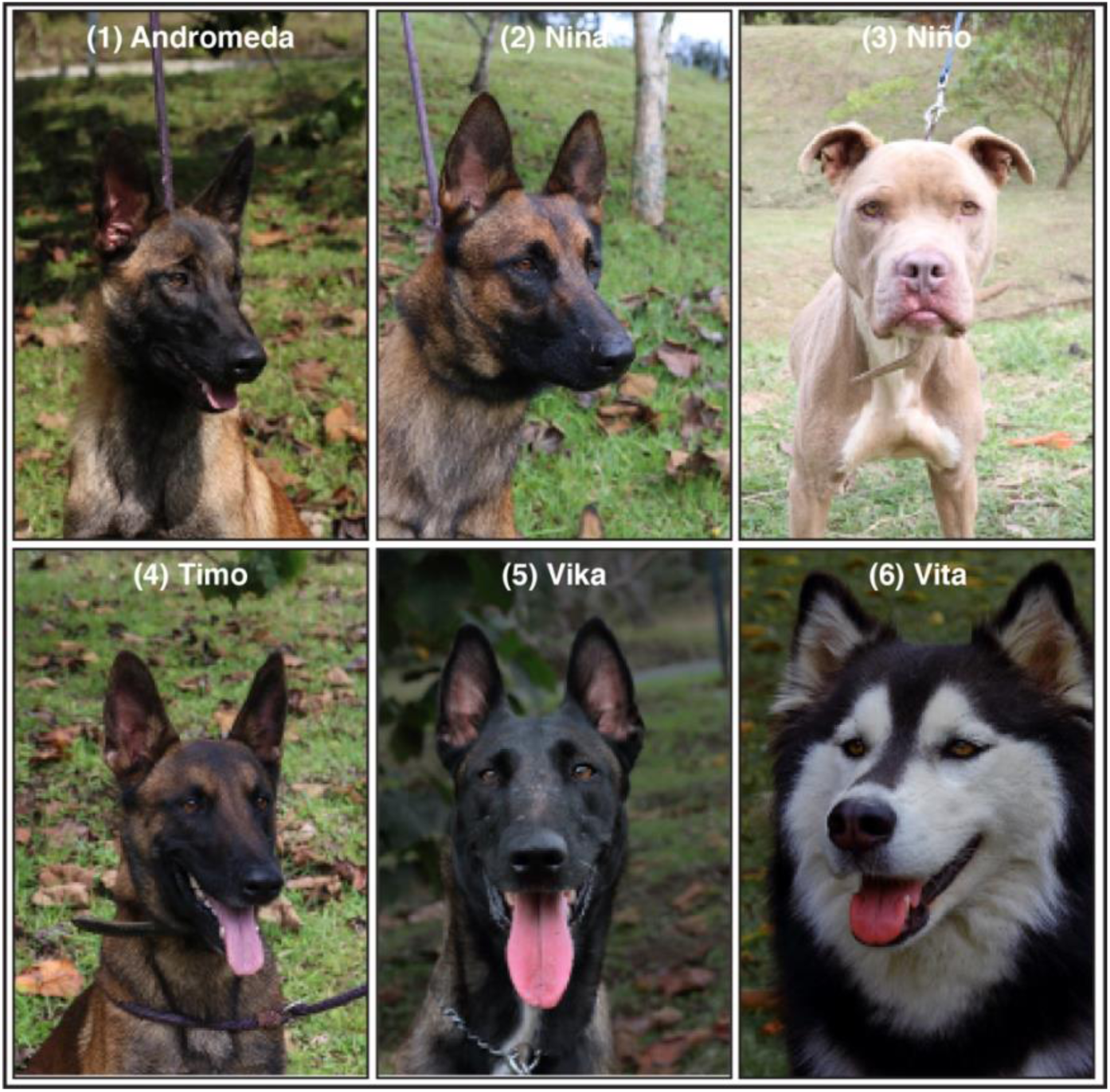
Pictures and identification of the six dogs trained for the scent-detection of SARS-CoV-2. (1) Andromeda, intact female, 6-mo, Belgian Malinois. (2) Nina, intact female, 25-mo, Belgian Malinois, (3) Niño, castrated male, unknown age, American Pit Bull Terrier. (4) Timo, intact male, 31-mo, Belgian Malinois. (5) Vika, intact female, 36-mo, Belgian Malinois. (6) Vita, intact female, 36-mo, first generation Alaskan Malamute x Siberian Husky.

The dog training component had three phases with a corresponding laboratory experimentation process aimed to generate data for statistical validation of our diagnostic test (see details in the Methods section). In the first phase (PI), called *in vitro* recognition, we trained our dogs to scent-detect SARS-CoV-2 in human respiratory secretions. Half of the positive specimens contained the active virus and the other half contained inactivated virus. We trained the dogs to identify the virus among negative controls made of sterile 0.9% saline solution. The specimens and the controls were presented in identical containers. For the second phase (PII), designated *in vitro* diagnosis, we used only active SARS-CoV-2 virus containers and, as distractors, saliva from one hundred healthy human subjects confirmed negative by rRT-PCR testing (see **Table 1**). In the third phase (PIII), *in vivo* diagnosis, the dogs learned to identify COVID-19 patients directly by smelling their bodies (work in progress).

**Table 1.**
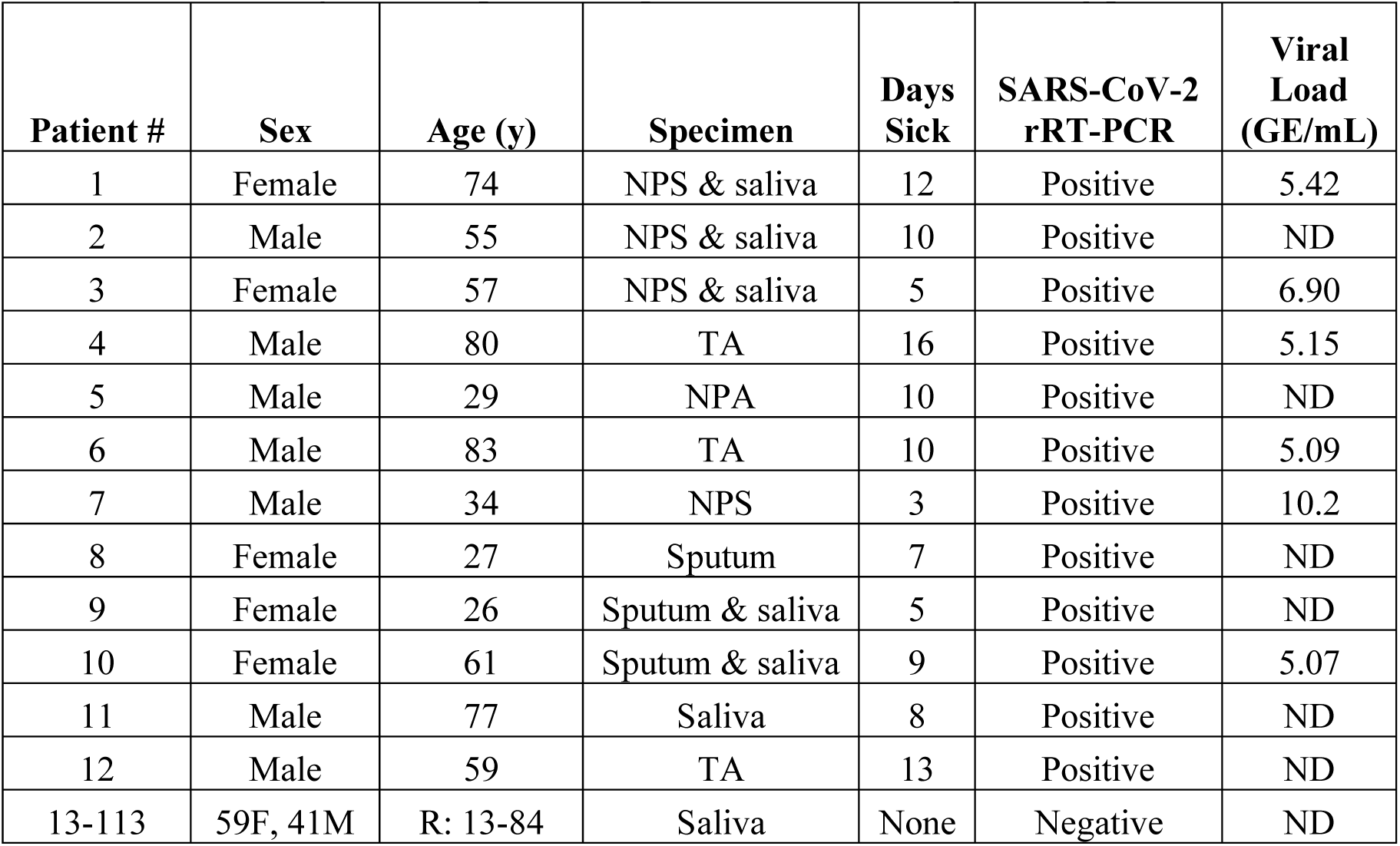
Human subjects who provided specimens for the dog training process.

It took 28 days to train all six dogs for *in vitro* recognition (PI) and 21 additional days to achieve *in vitro* diagnosis (PII). During the experimentation process in PI and PII, they scent-interrogated (i.e., to investigate by scent) 3,200 and 6,000 samples, respectively (N = 9,200). For each experiment, the dogs went through an open field arrangement of 10 x 10 samples (100) distanced 2 m in all directions (see supplemental **Movie 1**). We recorded and analyzed the data to determine *SEN, SPC, PPV, NPV, ACC, LR*, and the probability (*p*) of scent-detection by chance^15^ (See Methods).

### PI: *in vitro* recognition of SARS-CoV-2 in human respiratory secretions

We designed phase 1 to identify potential issues that would require additional work in phase 2 training. To do that, we subjected each dog only to the minimum number of experiments required and the trainers were not blinded in 75% of them (see **Table 2**). Nina and Vita scent-interrogated 1,000 samples each, Vika 700, Niño and Timo 200 samples each, and Andromeda 100 samples. The lowest individual performance was *SEN* 84.6% (Timo), *SPC* 95.8% (Vita), *PPV* 66.4% (Vita), *NPV* 98.8% (Nina), *ACC* 95.2% (Vita) and a *LR* of 21.3

**Table 2:**
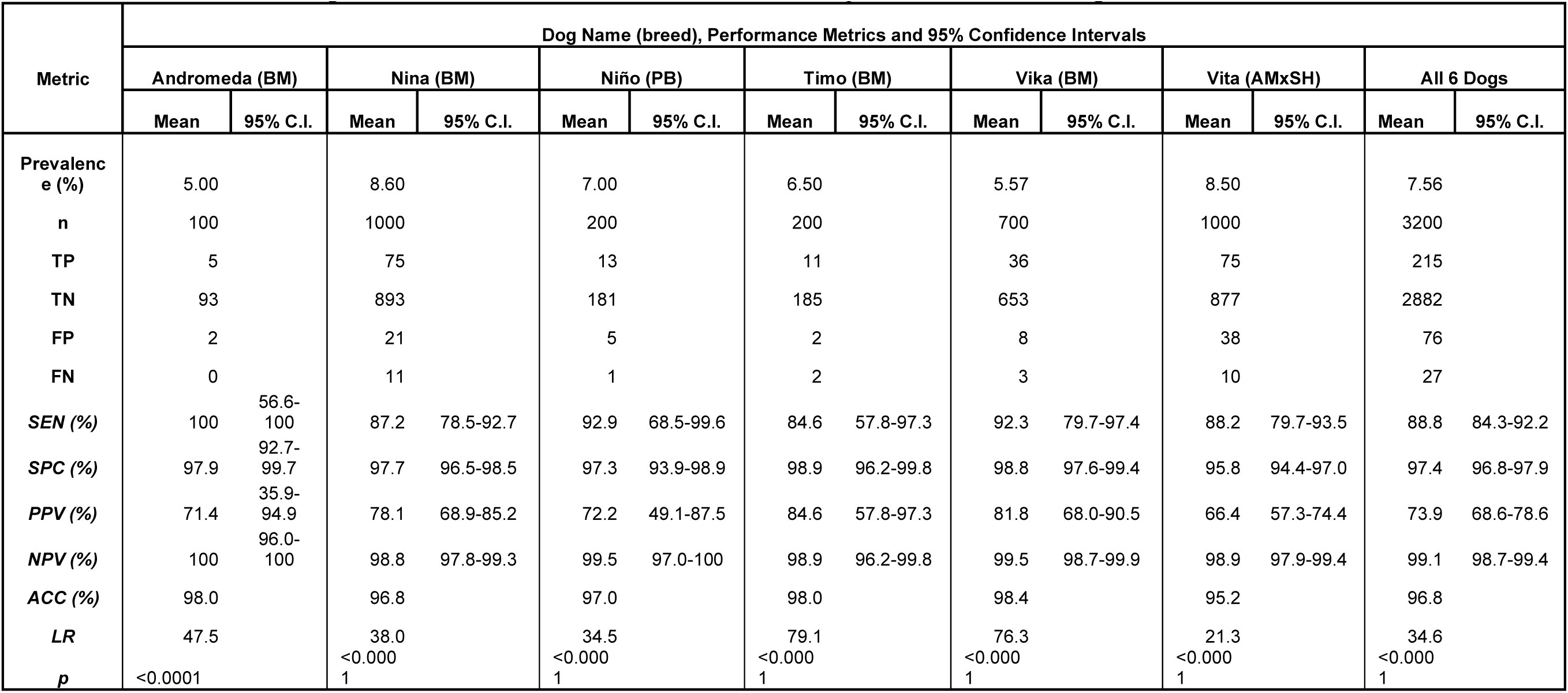
PI *in vitro* recognition. Performance metrics of each dog after four weeks of scent-detection training. In each experiment, the dogs have to interrogate 100 flasks containing sterile 0.9% saline solution or SARS-CoV-2. The position and prevalence of the virus was randomized for each dog. The trainers knew both variables for all experiments and interrogations.

(Vita). As a group, the six dogs achieved *SEN* 88.8% (95% CI 84.3 - 92.2), *SPC* 97.4% (96.8 - 97.9), *PPV* 73.9% (68.6 - 78.6), *NPV* 99.1% (98.7 - 99.4), *ACC* 96.8% and a *LR* of 34.6. The mean prevalence of positive samples for the PI experiments was 7.56% (range, 1%-10%).

During PI, we identified a couple of issues that needed special attention for the PII training. First, the relatively low *PPV* (73.9%) revealed a high rate of false positives in the early stages of training; 26 out of 100 patients diagnosed as “positive” by the dogs would not have COVID-19. These predictive values are proportional to the prevalence of the positive samples, i.e. when disease occurs at a lower prevalence, diagnostic tests will have lower *PPV* and higher *NPV*, and the opposite happens when disease prevalence is high^19^.

Therefore, we designed an experiment to determine if the *PPV* would improve by increasing the prevalence of positive samples to 20% without further training. In a field with 40 containers (8 with SARS-CoV-2 and 32 with saline), all six dogs identified the specimens with virus without a single mistake (i.e., no false positives or false negatives). Although a perfect performance is a desirable outcome, this experimental setting was not realistic because COVID-19 prevalence is currently lower than 1% in the human population. Therefore, during PII training, we worked the dogs with a narrower prevalence range (1% to 5%) and dissuaded their inclination to indicate false positives in order to receive more frequent rewards (see Methods). The second issue was the use of the inactive virus, which the dogs identified as well as the active virus. Dogs are not susceptible to infection by SARS-CoV-2 unless contact with a sick owner is intimate^20^, and, if infected, they do not develop disease^21^. However, it was mandatory for us to demonstrate that our virus enclosing device was safe for both dogs and trainers. During PII, we only used the active virus.

### PII: *in vitro* diagnosis of SARS-CoV-2 in human respiratory secretions

Phase 2 made a significant improvement in the metrics for every dog (see **Table 3** and **Figure 2**). As a group, they achieved *SEN* 95.5% (95% CI 90.4 - 97.9), *SPC* 99.6% (99.5 - 99.8), *PPV* 85.7% (79.2 - 90.5), *NPV* 99.9% (99.8 - 100), *ACC* 99.6% and a *LR* of 267. The *PPV* improved 12 percentile points, while the *NPV* was close to perfection, thereby suggesting an extremely low probability that any of our dogs would miss a positive case.

**Supplemental Movie 1:**
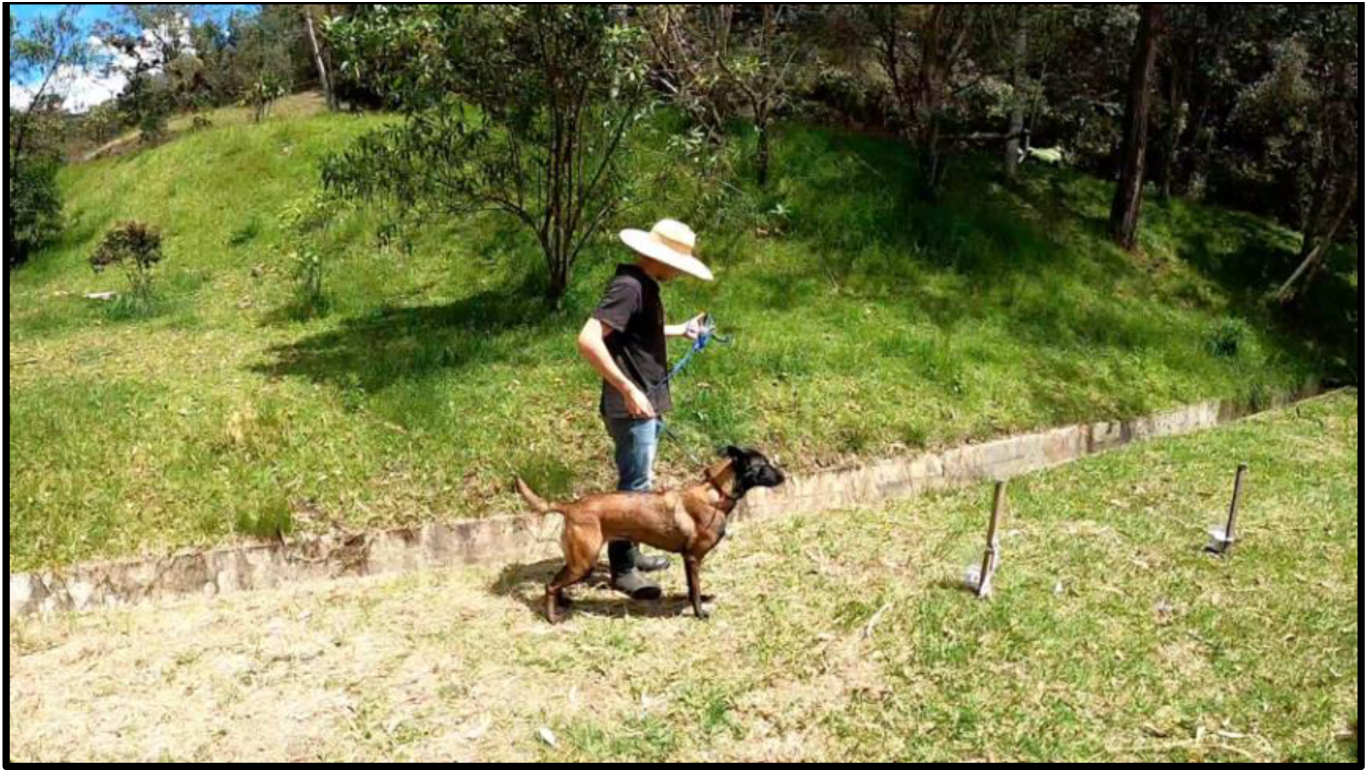
This video illustrates the experimental field as described in the text. Vika is displaying a perfect performance during the PII training.

**Table 3:**
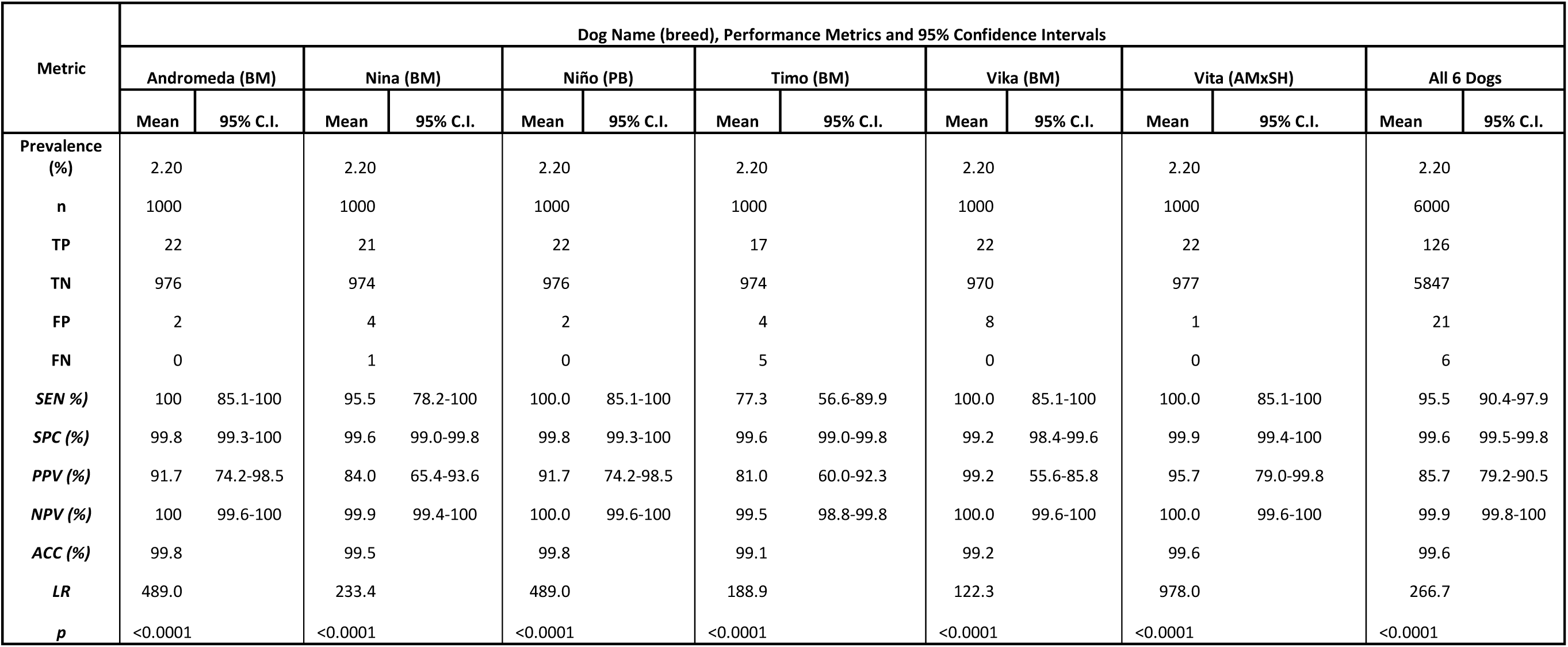
PII *in vitro* diagnosis. Performance metrics of each dog after seven weeks of scent-detection training. In each experiment, the dogs interrogate 100 flasks containing saliva from healthy human subjects or SARS-CoV-2 positive patients. The position and prevalence of the virus was randomized for each dog. The trainers were blinded on both variables for all 60 experiments.

**Figure 2:**
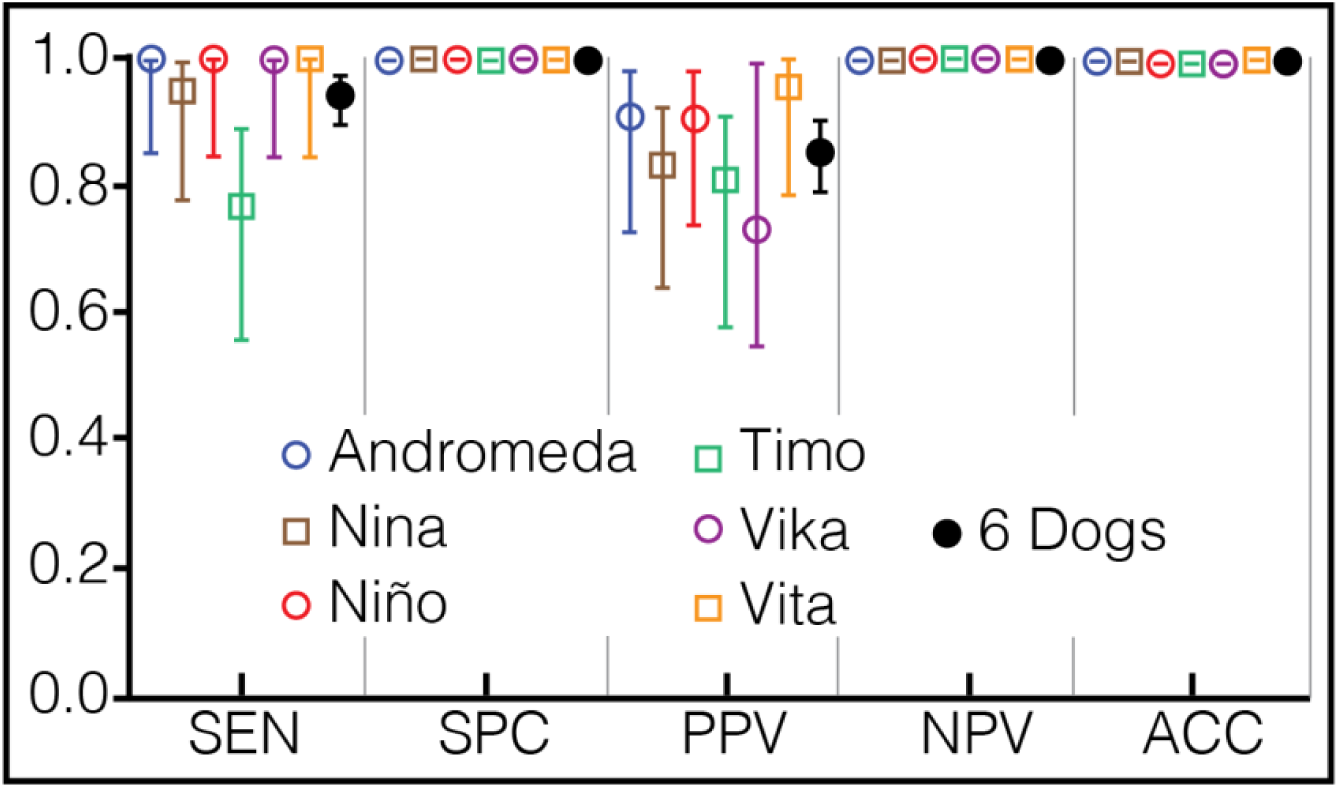
Phase 2: *in vitro* diagnosis. Each symbol has a different color to ease visualization of the dogs. The vertical lines above and below the symbols represent the 95% confidence interval for each metric.

### PII: limit of detection (LOD)

We determined the LOD using five freshly collected saliva specimens from COVID-19 patients. Half the specimen was destined for quantification of the viral loads in genome equivalents per milliliter (GE/mL), and the other half for serial 1:10 dilutions, making sure that the most dilute specimen of each patient had 0 log_10_ GE/mL. Then, we randomized the dilutions from each patient in our 100-set field and commanded every dog to search them. The results will be updated as we obtain them.

### PIII: in vivo diagnosis of SARS-CoV-2 in the body of patients with COVID-19

The results will be updated as we obtain them.

## Discussion

Accurate and timely detection of infected individuals is the most significant obstacle that societies have to overcome in order to contain SARS-CoV-2. Countries as diverse as South Korea, Iceland, Bahrein, and New Zealand, where the identification and prompt isolation of citizens harboring the virus independent of symptoms was prioritized, stopped the menacing pace of the pandemic within their territories, saving thousands of lives while maintaining healthy economies. On the contrary, countries where disease detection was not efficient, nor a priority, public health is undergoing devastating consequences, independent of their wealth and technological prowess. Our training system and dogs offer a reliable, accurate, and straightforward diagnosis of COVID-19 in seconds. As a unique feature, the dogs are immediately ready to attend the next patient only after a quick reward.

We tried to fulfill the ideal requirements relevant to validate a diagnostic test^22^. The 12 patients selected to scent-train the dogs provided a variety of sources, timing, and disease severity that included many of the real-life situations present in COVID-19 (Table 1). For respiratory specimens, we collected saliva, nasopharyngeal swabs or aspirates, and tracheal aspirates. Our subjects, males or females, offered wide ranges of age (26-83-yo), evolution at sampling time (3-16 days), symptomatology (mild, moderate, severe or lethal disease), and viral loads (5.07-10.2 log10 GE/mL). Despite all these features, we still need to include asymptomatic patients to guarantee that the dogs detect the whole spectrum of COVID-19 (work in progress).

Although breeds specialized in scent-detection are optimal for this task^23^, excellent olfaction is a genetic trait present in any healthy dog^24^. In our view, the general applicability of this project is broader if, besides being mentally and physically healthy, the only requirement for the dog is its inborn propensity to enjoy any activity involving its owner, independent of its trainability. We demonstrated that there was no significant difference in the performance of four Belgian Malinois, selected for more than a century for their trainability^25^, and a human-independent breed destined for sled work (Alaskan Malamute x Siberian Husky)^26^, or a fighting breed (pit bull) that, based on its history, has had little use for its nose^27^.

The potential applications of scent-detecting dogs to diagnose COVID-19 are limitless. For example, a diagnostic dog could be stationed in an airport to scent-interrogate all passengers upon check-in. The same principle could apply to ships, trains, factories, hospitals, malls, restaurants, stadiums, jails, or virtually any place where an immediate diagnosis could help prevent new infections. Furthermore, the possibility of training dogs for real-time diagnosis of many other infectious diseases may help humanity prepare for the next pandemic. We believe our method may aid the safe re-opening of economies and educational systems, while offering an efficient way to control the pandemic.

There are 135 million dogs with owners on the planet, and virtually all of them can smell an infection or certain cancers after appropriate training. If one million dog-owners accepted the challenge of scent-training (with inactive virus) to detect SARS-CoV-2, and each dog interrogated 800 people throughout their neighborhood (an average block in Manhattan harbors 1400 people), we could contain the pandemic in short order. The training process, although careful and methodical, is simple and easy to implement by anyone willing to follow a set of straightforward instructions that we would make available online after PIII. Their dogs could check families and neighbors, and the diagnosis of COVID-19 infection would be free and instantaneous for everyone. Self-isolation, simultaneously with rapid detection of contacts by the nearest dog, could be the best alternative to solve this problem because the needed technology is not available to most countries. It may not be feasible because there are many variables involved, and dogs learn to manipulate the trainer by signaling false positive specimens in order to get a reward. Some sort of accreditation system could be useful to make sure dog-owner duos are deployable. Local training groups and organizations could volunteer to help a lot if required. In the beginning, confirmatory testing by rRT-PCR would be needed until the public gain confidence in the method. Despite all the caveats, it might be worth a try.

## Methods

### Dog training

Using operant conditioning based on clicker-training and rewarding with food or prey-based play depending on the dog^28^, six animals of three different breeds were trained to detect the odor print of SARS-CoV-2 in saliva. Four Belgian Malinois aged six to 36 months, three females and one male (Andromeda, Nina, Timo, and Vika), one 36 month-old mixed Nordic female (Vita, first-generation Alaskan Malamute by Siberian Husky), and one male pit bull (Niño, unknown age) were trained and used to diagnose COVID-19. A dog rescue organization found the pit bull abandoned and tied to a tree. They castrated him and brought the dog to our training center for rehabilitation due to extremely aggressive behavior. Before starting this project, we had trained the dogs in basic obedience, and had the pit bull fully rehabilitated.

Respiratory secretions from 12 patients with COVID-19 pneumonia (rRT-PCR positive for SARS-CoV-2), and saliva from 100 healthy volunteers were obtained (Table 1). For every experiment, the position of the samples (1 to 100) and the prevalence (1% to 10%) of the virus specimens within the negative controls were randomized with a mobile phone. During training, the total number of containers (n), the proportion of positive samples (prevalence), and their position in the sample line were modified, from simple to complicated patterns, thereby creating different scent problems for the dogs to solve. The limit of viral detection (GE/mL) for each dog was determined. Also, to discard the possibility of infection during the study, saliva samples from dogs and trainers were tested by rRT-PCR at the end of the second and third phases.

### Experimentation after scent-detection training

After finishing scent-training, the interrogation setup, to determine the accuracy of our dogs to detect SARS-CoV-2, consisted of 100 wood sticks standing up 80 cm above the ground on a grass field. The setup was large enough to accommodate ten rows and ten columns separated two meters from each other in all directions. Three kinds of 2-mL specimens were prepared, under a biosafety class II laminar flow cabinet, using 212 sterile, scent-free flasks. One-hundred flasks had 0.9% sterile saline solution, 100 had rRT-PCR-negative saliva, and 12 flasks had COVID-19 positive respiratory secretions. The saliva and respiratory secretions specimens were diluted (1:1 volume) in 0.9% sterile saline solution to preserve the virus^29^, without altering the odor of the specimen. The recipients were brand-new, 130-mL transparent glass flasks with a metallic mouth (4.5 cm diameter). They were covered with a 10 x10 cm^2^ piece of DuPont Tychem™ and sealed hermetically with a screwable metallic cap that had a 1 cm hole in the center. Therefore, the dogs were allowed to smell the VOC while preventing any exposure to the virus. We confirmed that this contraption prevented exposure by subjecting dogs and trainers to rRT-PCR for SARS-CoV-2 in nasopharyngeal secretions at the end of the experimental process (all were negative). Saliva, for the negative controls, was collected from 100 asymptomatic volunteers; on the other hand, the positive specimens were from 12 hospitalized COVID-19 patients whose diagnosis had been demonstrated by rRT-PCR (Table 1). All the specimens were stored in different refrigerators at 4°C.

During experimentation, the trainers were blinded regarding the position and number of positive specimens. On command, the dog would scent-interrogate flasks 1-10 (first row), return searching flasks 11-20 (second row), and continued in a zig-zag pattern until reaching the last flask (tenth row). Between rows, the trainer stimulated his dog for a few seconds offering a tidbit and playing with a ball or a tug. Each time the dog correctly marked SARS-CoV-2 by downing in front of the positive specimen, a clicker sound announced it, and the trainer would immediately reward his dog. The number of positive specimens in the 100-flask field marked the prevalence of COVID-19 per experiment: i.e. 1%-10% in PI, 1%-5% in PII. A mobile phone app was used to randomize the position and prevalence of positive specimens. The experimental process in PII was filmed and will be available online. The data was collected in a 2 x 2 contingency table and processed to calculate *SEN, SPC, PPV, NPV, ACC*, and *LR*^15^. We applied the Fisher’s Exact Test to determine the 95% confidence interval of each of the first four metrics and to challenge the null hypothesis (i.e., that the dogs found the positive samples by chance) because the dogs rarely made mistakes (i.e., false negatives or false positives), thereby generating 0 to 5 values in the respective cells of the contingency table for almost all the experiments.

### rRT-PCR assay and RNA quantification

The SARS-CoV-2 molecular diagnosis was done at the Genomic One Health Laboratory (Colombia\Wisconsin One Health Consortium) at the Universidad Nacional de Colombia. Canine nasal and oropharyngeal swabs and human nasopharyngeal aspirates were used. The RNA extraction was carried out using the ZR viral extraction kit (Zymo Research) from 140-μL of specimens. Instructions provided by the manufacturer were followed and the sample was eluted into 20 μL. The CDC 2019-Novel Coronavirus Real-Time RT-PCR Diagnostic Panel (Integrated DNA Technologies)^30^ and Berlin-Charité E gene protocol for SARS-CoV-2^31^ were used to detect virus nucleocapsid (N1 and N2) and Envelope genes respectively. All rRT-PCR testing was done using Superscript III One-Step RT-PCR System with Platinum Taq Polymerase (Thermo Fisher Scientific). Each 25-μL reaction contained 12.5 μL of the reaction mix, 1 μL of enzyme mix, 0.5 μL of 5 μmol/L probe, 0.5 μL each of 20 μmol/L forward and reverse primers, 3.5 μL of nuclease-free water, and 5 μL of RNA. The amplification was done on an Applied Biosystems 7500 Fast Real-Time PCR Instrument (Thermo Fisher Scientific). Thermocycling conditions consisted of 15 min at 50°C for reverse transcription, 2 min at 94°C for activation of the Taq polymerase, and 40 cycles of 3 s at 94°C and 30 s at 55°C (N gene) or 58°C (R gene), and 3 min at 68°C for the final extension. SARS-CoV-2 assays were simultaneously ran along with internal control genes for canine (glyceraldehyde-3-phosphate dehydrogenase-GAPDH^32^) and human specimens (Ribonuclease P-RP^30^) to monitor nucleic acid extraction, sample quality, and presence of PCR reaction inhibitors^33^. To monitor assay performance, positive template controls and no-template controls we also incorporated in all runs. Biosafety precautions were followed during the work-flow to minimize PCR contamination. For rRT-PCR qualitative detection, a threshold was set in the middle of the exponential amplification phase of the amplification results and a specimen was determined as positive for SARS-CoV-2 when all controls exhibited expected performance and assay amplification fluorescent curves crossed the threshold within 40 cycles (Ct <40). For rRT-PCR quantitative detection on human specimens, an analysis of copy number and linear regression of the RNA standard was used.

### Preparation of *in vitro* RNA Transcript as standard

An *in vitro* RNA transcript of the SARS-CoV2 envelope gene was generated as a standard for rRT-PCR quantitative detection on human specimens. Viral RNA from a positive clinical sample was used as initial template for *in vitro* RNA transcription. cDNA was synthetized using SuperScript™ III First-Strand Synthesis System and random hexamers primer (Thermo Fisher, USA).

Double-stranded DNA containing the 5′-T7 RNA polymerase promoter sequence for the SARS-CoV-2 complete E gene sequence, was obtained using DreamTaq Hot Start PCR Master Mix (Thermo fisher, USA) and E-Std-T7-Fwd (TAA TAC GAC TCA CTA TAG GGG CGT GCC TTT GTA AGC ACA A), and the E-Std-Rev (GGC AGG TCC TTG ATG TCA CA) primers^34^. The DNA was finally transcribed using the MEGAscript T7 Transcription Kit (Thermo Fisher Scientific). The RNA transcripts were purified with Ampure XP beads (Belckman Counter, USA) and quantified with a Qubit fluorometer by using a Qubit RNA HS Assay Kit (Thermo Fisher Scientific). All commercial reagents were used following manufacturer instructions.

### Assay efficiency and analytical sensitivity (LOD)

The *in vitro* RNA transcript standard was used to assess LOD and assay efficiency using a standard curve. Serial 10-fold dilutions of quantified *in vitro* RNA transcript were prepared in triplicates per dilution. The LOD for each assay was defined as the highest dilution of the transcript at which all replicates were positive. The efficiency (E) was estimated by linear regression of the standard curve using the equation^35^ (E) = [10 _(1/slope)_] – 1. The LOD and E of the SARS-CoV-2 assay were determined to warrant consistency with what has been previously demonstrated^36^. The intra- and inter-assay variability were also calculated using the in vitro RNA standard. To assess intra-assay variation, the RNA standard was used at 2 and 6 log_10_ copies/reaction by triplicate in a single assay. To assess inter-assay variation, the RNA standard was tested at 2 and 6 log_10_ copies/reaction by triplicate in two separate PCR assays. Mean, standard deviation, the coefficient of variation of the cycle threshold (Ct) and copy numbers were also determined.

## Acknowledgments

The authors and our society are genuinely in debt to our economic supporter, Dr. Mauricio Palacio. His generous donation allowed us to do this research. The authors also wish to thank the Grupo ISA for their donation, which included the RNA extraction kit and reagents for the Berlin PCR Amplification. We also want to recognize master trainer and breeder Christoph Joris, because thanks to his methods, his teaching, and his example, we were able to accomplish this project. We thank sincerely, Dr. Tonie E. Rocke, for her insightful comments to improve so significantly the quality of the manuscript.

## Authors Contributions

OV came up with the idea, obtained the funding, designed and supervised the training and the experimental process, and wrote the manuscript.

AFV directed the training process and supervised the dog experiments in the field.

AM, FO and EO trained the dogs and participated in all the experiments with them.

AFV, AM and FO obtained and processed the saliva specimens from 100 healthy volunteers.

MA coordinated, obtained, processed, made available for experimentation, and took medical care of the patients with COVID-19 who provided the respiratory specimens to train our dogs.

KC, LP, AC and YA performed the molecular biology work.

JMG and JCC took care of the patients with COVID-19 who provided the specimens to train our dogs.

YA directed the logistics and organization of the experimental process with COVID-19 and control patients within *Hospital Universitario San Vicente Fundación*.

JPH-O and JO supervised the molecular biology work, supported the project with research funds, read the manuscript, and made invaluable comments to improve its quality.

## Ethical Statement: animal usage and human samples

We did not subject our dogs to any kind of pressure for training. We did not starve the dogs and did not need to make them fanatics for food, toys, games, or anything else. Since it is impossible to force a dog to do scent-work, our methods are exclusively positive, rewarding every correct response, and being indifferent to any mistake. All human subjects read and signed their informed consent.

